# Clustering scRNA-seq data via qualitative and quantitative analysis

**DOI:** 10.1101/2023.03.25.534232

**Authors:** Di Li, Qinglin Mei, Guojun Li

## Abstract

Single-cell RNA sequencing (scRNA-seq) technologies have been driving the development of algorithms of clustering heterogeneous cells. We introduce a novel clustering algorithm scQA, which can effectively and efficiently recognize different cell types via qualitative and quantitative analysis. It iteratively extracts quasi-trend-preserved genes to conform a consensus by representing expression patterns with dropouts qualitatively and quantitatively, and, then automatically clusters cells using a new label propagation strategy without specifying the number of cell types in advance. Validated on 20 public scRNA-seq datasets, scQA consistently outperformed 9 salient tools in both accuracy and efficiency across 16 out of 20 datasets tested, and ranked top 2 or 3 across the other 4 datasets. Furthermore, we demonstrate scQA can extract informative genes in both perspectives of biology and data wise by performing consensus, allowing genes used for landmark construction multiple characteristics, which is essential for clustering cells accurately. Overall, scQA could be a useful tool for discovery of cell types that can be integrated into general scRNA-seq analyses.

---

Advances of single cell RNA sequencing (scRNA-seq) technologies have led to a fresh look at development and cellular heterogeneity of transcriptomes at single-cell resolution and enhanced to understand how cells make up complex and changing system Shalek et al. (2013); (Buettner et al. 2015). With the sequencing cost becoming cheaper, vast amounts of scRNA-seq data of remarkably different scale and accuracy (Ziegenhain et al. 2017; Ding et al. 2020; Mereu et al. 2020) have been produced. Many researchers have been devoting to unveiling the biological meaning by analyzing the scRNA-seq data, but analyses remain immensely challenging (Kiselev et al. 2019) due to high rate of noises and high dimensionality of the data.

One common phenomenon of scRNA-seq data from different protocols is so called “dropout”, describing zero expression for genes that show non-zero expression in only a proportion of cells (Kharchenko et al. 2014). Several reasons can be found to response for dropout events, including technical artifacts such as amplification bias and low sequencing depth, resulting in zero expression measurements of genes (Jiang et al. 2020). Beyond that, biological factors should also be account for this phenomenon. For many years, plenty of methods have been considered to address high sparseness as the result of dropouts. Principal component analysis (PCA) (Wold et al. 1987) and other dimension reduction strategies are often used to combine clustering methods for identifying cell types. For instance, SC3 (Kiselev et al. 2017) is a consensus method applying *k*-means on transformed distance matrices using PCA or eigenvectors of Laplacian, and eventually classifies cells by hierarchical clustering strategy. Seurat (Satija et al. 2015) adopts shared nearest neighbor (SNN) graphs and Louvain community detection after performing PCA. RaceID2 (Grun et al. 2016) is designed for the identification of rare and abundant cell types using *k*-medoids based on Pearson correlation matrix. Another popular method, CIDR (Lin et al. 2017) uses PCA by considering dropout events to improve similarities. Nowadays, the application of deep learning to scRNA-seq data analysis is rapidly evolving, such as scGNN (Wang et al. 2021), which integrates graph convolutional network into multi-modal autoencoders for gene imputation and cell clustering. Apart from introducing dimension reduction strategies to mitigate the impact of high dimensionality and sparseness, there are also special imputation methods to substitute zeros with numerical values by fitting data with a moderate model, such as SAVER (Huang et al. 2018), MAGIC (van Dijk et al. 2018), ScImpute (Li and Li 2018) and SCDD (Liu et al. 2022).

Recently, a different view of dropouts has been adopted from the literatures that supports that the dropouts could be of the essence. Qiu (Qiu 2020) clusters cells based merely on binary information, with the results being compatible with other conventional methods. M3Drop (Andrews and Hemberg 2019) employs genes with higher dropout rate as useful features for downstream analyses. scBFA (Li and Quon 2019) shows that the features obtained by applying binary PCA to the expression data can be used to cluster cells accurately. In consideration of the general thought of treating dropouts as noises, Kim (Kim et al. 2020) points out that imputation or normalization before resolving the cellular heterogeneity may lead to inevitable loss of biological signals for unique molecular identifiers-based (UMI-based) data. Meanwhile, it has been widely evidenced that technical zeros can be decreased in UMI-based data while modeling zeros using different methods that may lead to wide disagreement for downstream analyses (Silverman et al. 2020; Svensson 2020; Sarkar and Stephens 2021). Hence, choosing an approach to correct zeros should be scrupulous since the simple way to treat dropouts or zero expressions as noises could be damaged to original data.

Inspired by what mentioned above, we propose a new algorithm scQA (an architecture for clustering **S**ingle-**C**ell RNA-seq data based on **Q**ualitative and **Q**uantitative **A**nalysis), which can effectively and efficiently cluster cells at various scale based on so called landmarks and each indicates the consensus of genes with similar expression patterns. In our opinion, treating dropouts as some sort of signals filling with noises, and meanwhile, combining some genes with similar qualitative expression vectors together can reduce noises to highlight signals. For this kind of qualitative trend-preserved (Supplemental Method 1) genes, their quantitative expressions should be considered since two closely related subpopulations may exhibit the same dropout patterns with very different levels of expression. In consequence, scQA adopts a novel feature extraction method which constructs the consensus vector of genes whose qualitative and quantitative expressions under certain cells are of similar trend, which we called quasi-trend-preserved genes (Supplemental Method 1). Differing from other dimensionality reduction methods, scQA not only keeps as much information that both qualitative and quantitative patterns contain as possible, but also reduces the noises by performing consensus. Subsequently, two landmark matrices generated by Landmark Constructor (LC_1_ and LC_2_) are combined into Cluster Constructor (CC) for clustering cells by a novel label propagation strategy, whose initial labels are given by generating seeds. The numbers of landmarks and cell clusters identified by scQA are automatically determined. Tested on 20 public scRNA-seq datasets (the number of cells ranges from 56 to 26,830) in which cell types were well-annotated, scQA substantially outperformed other salient tools compared in this paper in a time efficient manner. In addition, we illustrated that scQA can capture similar genes forming a landmark and can accurately cluster cells by multiple different landmarks. We also demonstrated that genes forming a landmark can have both the characteristics of differentially expressed genes associated with cell types, hub genes derived by constructing a weighted gene co-expression network, and marker genes of biological significance.

## Results

### Overview of scQA

scQA mainly consists of three modules: qualitative landmarks constructor (LC_1_), quantitative landmarks constructor (LC_2_), and clusters constructor (CC). The first two modules are to extract features in genes by generating consensus expression pattern for every cluster of quasi-trend-preserved genes that is referred to as a landmark. In this manner, signals are naturally captured while noises are eliminated. The third module is to cluster cells by a label propagation strategy.

The module LC_1_ takes as input the binarized gene expression matrix after pre-processing which performs filtration, normalization and picking up variable genes. Initially, for each pair of genes, we calculate a similarity score based on their binary expression values under all cells, and genes with their pairwise scores beyond a prespecified threshold are grouped to form initial clusters as a prior information which will be used to reduce the impact of order of the genes processed by LC_1_. Then other genes will be added into a single existing cluster or isolated from the current clusters as a new one based on its expression pattern. Removing those less significant resulting clusters, we then generate a consensus pattern as a landmark for each remaining cluster of genes of qualitative quasi-trend-preserving pattern. These landmarks form a low-dimensional expression profile of the cells, the first constructed cell-landmark matrix, which will be used to generate seeds in CC.

Taking the pre-processed gene expression matrix as input, LC_2_ first partitions all genes into a few groups by employing a binning strategy, and then clusters genes in each group such that genes in the same cluster are of similar expression patterns. As did in LC_1_, removing those less significant resulting clusters, we then generate a consensus as a landmark for each remaining cluster of genes of quantitative quasi-trend-preserving pattern, forming the second cell-landmark matrix, which will be used to create a directed *k*-nearest neighbor graph in CC.

The module CC starts with a directed *k*-nearest neighbor graph constructed with nodes representing cells and an edge arrowed from node *i* to node *j*, we say that node *j* is a neighbor of node *i*, if cell *j* is ranked at top *k* in similarity to cell i. Then clustering cells can be achieved in four steps. In the first step, for each node, we assign a score to the node the average similarity between the node and its neighbors. We then determine a distribution of seeds according to the scores defined on individual nodes (see “Methods” section for details). In the second step, we label each seed with an integer number, and then propagate each label from the labeled seed to the unlabeled nodes each either is a neighbor of the seed or of the seed as its neighbor, and no less similar to the seed than to other seeds. Now most of nodes are labeled and each cluster is created by the nodes of same label. In the third step, we remove the clusters of size less than a prespecified threshold and merge similar clusters to reduce complementary entropy (Supplemental Method 2). In the last step, we add an unlabeled node to a cluster and label it if the number of neighbors in the cluster of the unlabeled node minus the average number of neighbors of each node in the cluster is greater than zero and maximized among the current clusters, otherwise, we establish a new cluster for the unlabeled node to be added to and with a new label assigned. Then we iterate the label propagation process until each node label no longer changes.

### scQA effectively and efficiently clusters cells

To test scQA, we compared it with nine salient algorithms, including SC3, Seurat, CIDR, pcaReduce (Zurauskiene and Yau 2016), SIMLR (Wang et al. 2017), TSCAN (Ji and Ji 2016), RaceID2, scGNN and scHFC (Wang et al. 2022) (see “Methods” section) on twenty public datasets with cells ranging from 56 to 26,830. The details of the nine algorithms and the twenty datasets are described in Supplemental Table S1 and S2. To systematically evaluate how well the competitive tools recover the true cell types, we calculated Adjusted Rand Index (ARI) (Hubert and Arabie 1985), Fowlkes-Mallows index (FMI) (Fowlkes and Mallows 1983), Normalized Mutual Information (NMI) (Vinh et al. 2010) and Jaccard index (JI) (Jaccard 1912) by which we further did comparisons with results shown in Fig. 2, Supplemental Fig. S1, Fig. S2 and Fig. S3 respectively. As shown in Fig. 2a, scQA outperformed all compared algorithms on sixteen datasets, and competitive with Seurat, scHFC and CIDR which respectively got the highest ARI score on one of the rest four datasets (Fig. 2b). Especially, scQA was significantly superior to all the others on Yan, Deng, Xin, Muraro, Segerstolpe, Hermann, Romanov, Zeisel, Campbell and Shekhar datasets with ARI score over 10% higher than the second-best algorithm and even more than 30% higher on Xin dataset (Fig. 2b). On average, scQA achieved ARI score of 0.71 (Supplemental Table S3), which surpassed second place (scHFC) by 17%. Supplemental Figures S1, S2 and S3 showed similar comparison results in NMI, FMI and JI scores. scQA improved on runner-up RaceID2 by 18% in JI score and on the runner-up scHFC by over 15% in FMI score, while lagged behind the best tool by lower 1% in NMI score (Supplemental Tables S4-S6).

**Figure 1.**
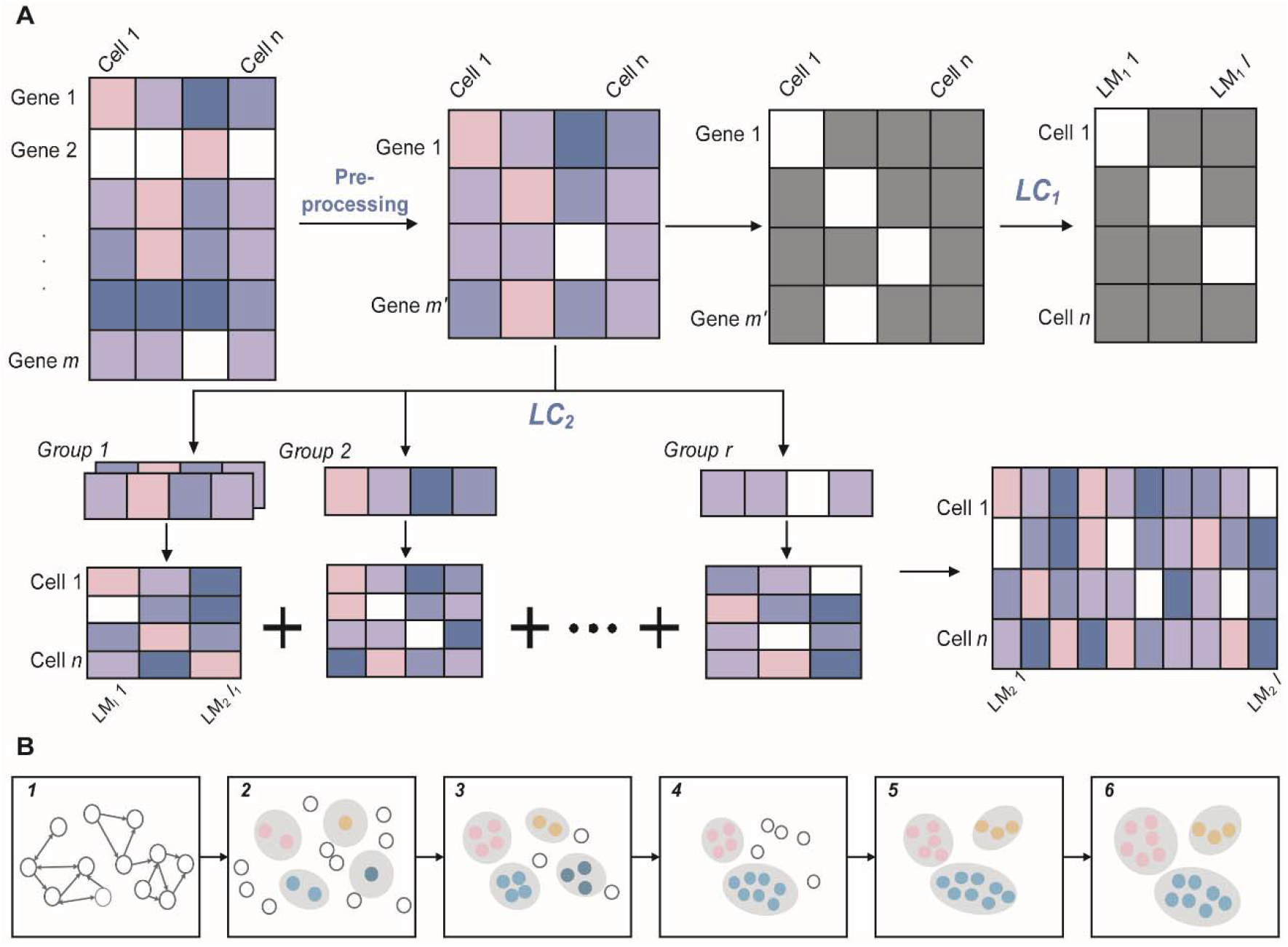
Illustration of scQA. (*A*) The raw gene expression matrix is processed by filtration, log-transformed and highly variable genes (HVGs) are retained. The matrix after pre-processing is fed to LC_1_ and LC_2_ to extract consensus features respectively. LC_1_ binarizes the matrix first and persists to cluster genes whose expression patterns are of qualitative quasi-trend-preserved. A qualitative cell-landmark matrix is generated in LC_1_. Genes before sending to LC_2_ need to be roughly grouped according to their expression patterns. Then LC_2_ clusters genes in each group such that each cluster consists of quantitative quasi-trend-preserved genes. The two cell-landmark matrices on behave of the two low-dimensional representations are used to cluster cells. (*B*) Illustration of clustering cells. In step 1, a directed *k*-nearest neighbor graph is created in terms of quantitative cell-landmark matrix. In step 2, vertices with high scores are regarded as seed candidates of which similar ones are merged into a single one. Each seed is assigned a unique label which is propagated to form a cluster in step 3. In step 4, clusters are screened and merged to obtain the highly reliable clusters. All unlabeled vertices are either added to the current clusters or isolated from the current clusters to form new ones in step 5. In the last step, label propagation process is iterated to output stable solution.

**Figure 2.**
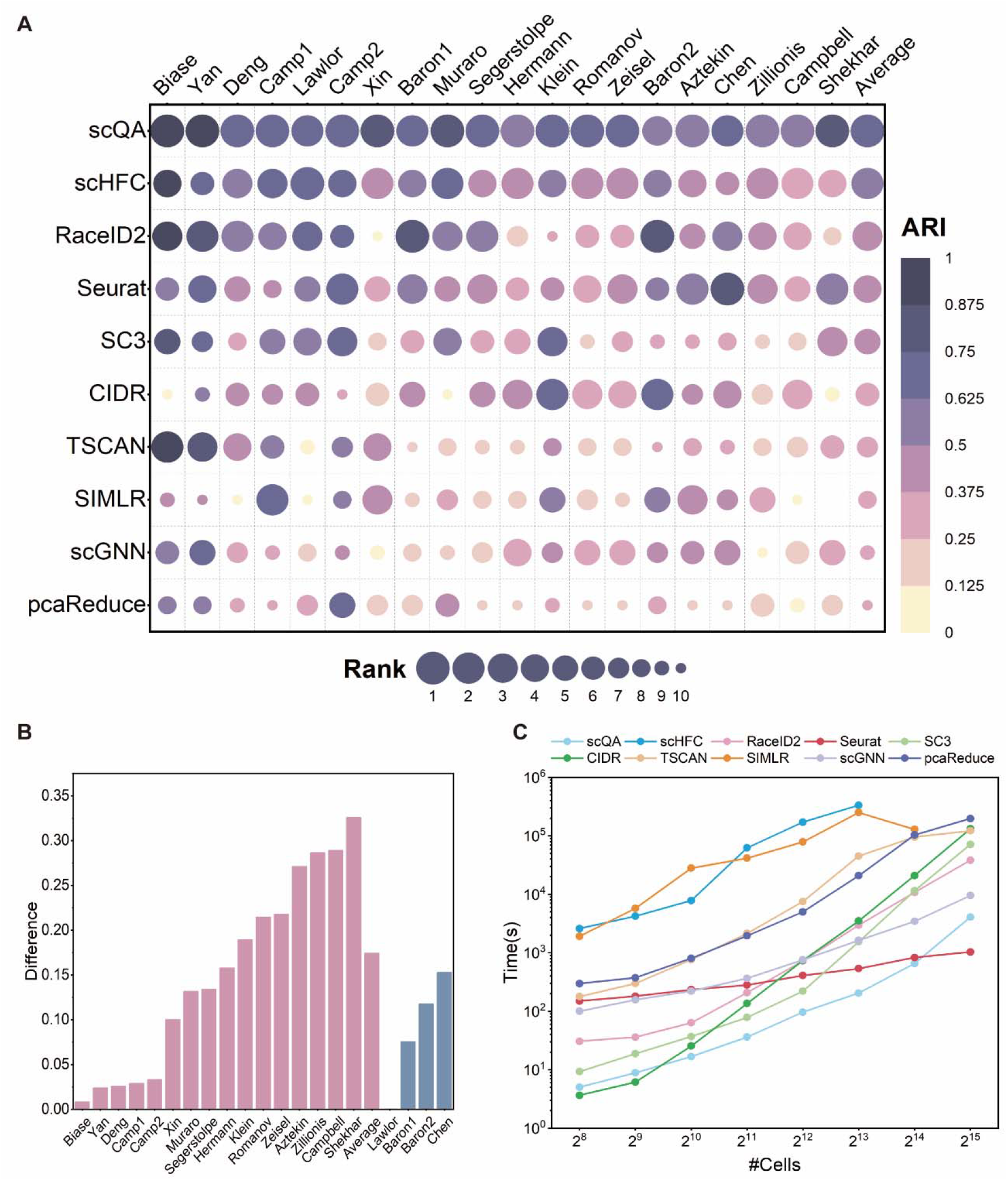
Evaluation of the compared algorithms. (*A*) Comparison of distributions of ARI scores against twenty datasets of the ten compared algorithms. Names of the datasets are represented on the *x*-axis, and methods are shown on the y-axis. Each scatter point reflects the clustering performance with color representing the ARI score and size indicating the rank. The vacancy in the plot indicates that SIMLR failed to estimate the number of clusters and cluster cells on Shekhar dataset due to excessive memory usage. Average ARI for all datasets except the vacancy is displayed in the last column. (*B*) Difference of ARI between scQA and the second-best methods on 16 datasets where scQA achieved the highest ARI (pink bars) and difference of ARI between the best methods and scQA on 4 datasets where scQA failed to achieve the highest ARI (blue bars). (C) Comparison of distributions of running times against the number of cells. SIMLR failed to estimate the number of clusters and cluster cells on the dataset with 2^15^ cells due to excessive memory usage. Running times over 10^6^ s were not displayed.

We evaluated their efficiency by comparing their distributions of running times against the numbers of cells with fixed number of genes (Fig. 2c). To test it, we generated datasets by varying the number of cells from 2^8^ to 2^15^ with number of genes fixed to 20,000 using a human liver immune scRNA-seq dataset (Zhao et al. 2020). All parameters of algorithms were set as before (see “Methods” section). Since some methods utilize multiple CPU cores for parallel computing whereas others use only a single thread to perform clustering, we compared the amount of CPU time spent for each method and all experiments were carried out on a Linux server. We can see that CIDR is the fastest method when number of cells is less than 1,000, and scQA is significantly and consistently efficient than other methods for number of cells less than 2^15^. However, when the number of cells reaches 30,000, scQA is slightly slower than Seurat with both time orders of magnitude being 10^3^. The methods scHFC and SIMLR are the two slowest methods compared to others whereas scHFC’s running time exceeded 10^6^s as the number of cells went beyond 2^14^, and SIMLR ran more efficient as the number of cells went beyond 2^13^ via using its second strategy for large-scale datasets. In general, scQA is faster than other top-performing methods and even as the number of cells exceeds 30,000, it is still acceptable.

The cell clustering results of scQA were illustrated by 2D visualization of embedded representations using uniform manifold approximation and projection (UMAP) (Fig. 3) and t-distributed stochastic neighbor embedding (t-SNE) (Supplemental Fig. S4). We displayed three datasets, labeled with the original labels (Fig. 3a-3c) and clustering labels (Fig. 3d-3f) respectively. In Camp1 dataset (Fig. 3a, 3d), scQA identified five clusters labeled with original annotations except that part of neuron cells were mistakenly assigned to dosal cortex neuron cluster. In the visualization plot, one can see that these two clusters are indeed intertwine in the raw data. In Camp2 dataset (Fig. 3b, 3e), six of seven clusters were recognized, and the cluster labeled with endothelial was divided into two separate clusters, since the UMAP plot showed the two parts of endothelial cells being relatively far away. In addition, mature hepatocytes and immature hepatocytes are close to hepatic endoderm cells in the UMAP plot and incorrectly recognized as the same cluster in scQA. A similar phenomenon occurred in Muraro dataset. As shown in Fig. 3c and Fig. 3f, ductal cells and acinar cells are close enough to each other on the original data, clustered into one group in scQA. However, the two cell types, endothelial cells and mesenchymal cells are far apart in the original data but incorrectly grouped into one cluster when performing clustering analysis. One possible reason for this is that the number of cells belonging to endothelial cell type is extremely small to be ignored during clustering and then was grouped into the cluster of mesenchymal cells by re-clustering.

**Figure 3.**
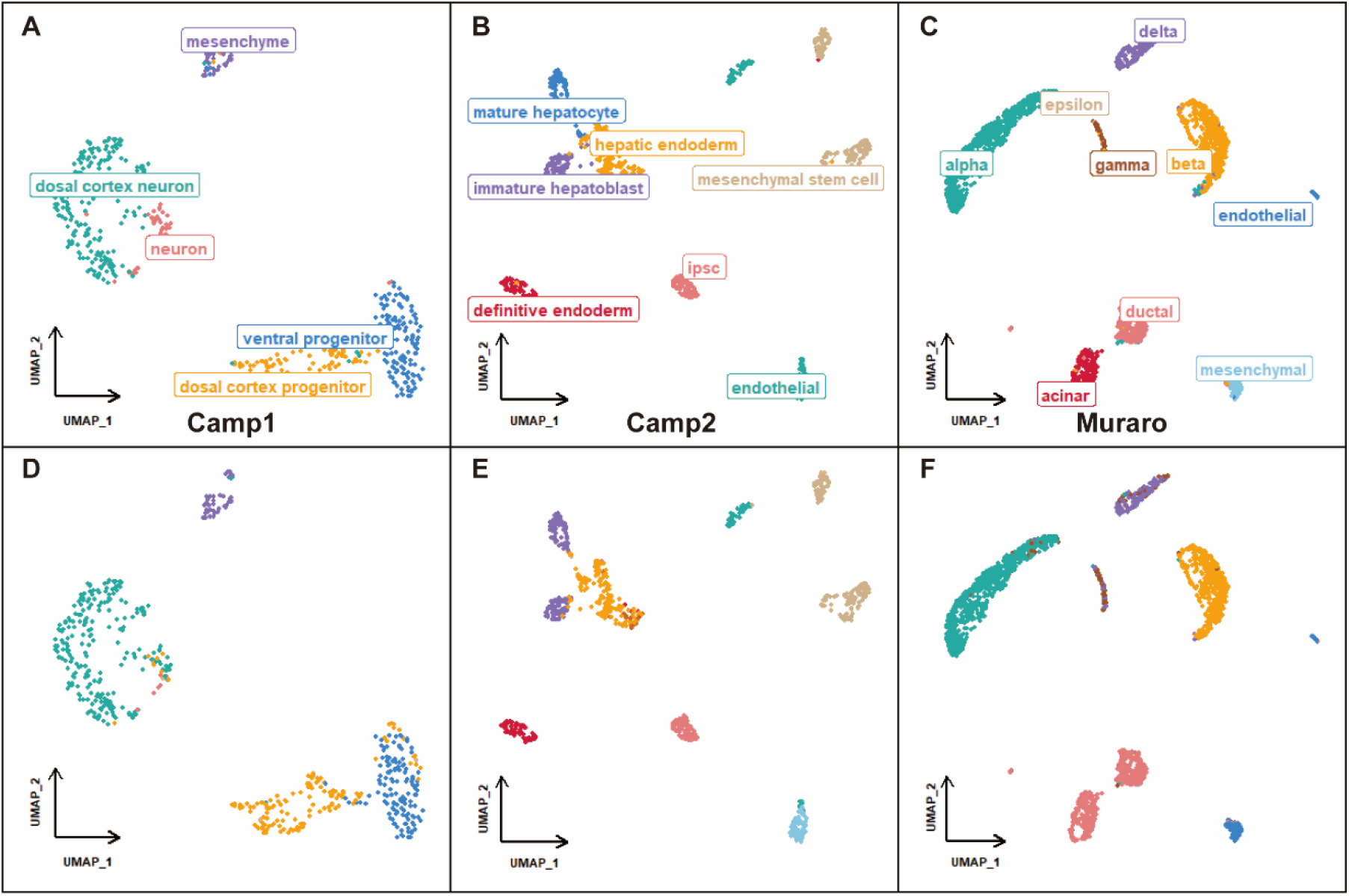
UMAP visualizations on three datasets labeled with original labels and cluster labels respectively. (*A-C*) UMAP of Camp1 dataset, Camp2 dataset and Muraro dataset with original labels. (*D-F*) UMAP of Camp1 dataset, Camp2 dataset and Muraro dataset with cluster labels.

### Self-reliability of landmarks

As we have demonstrated that scQA can accurately identify distinct cell types, we proceed to analyze the characteristics of the identified landmarks both internally and externally. In this section we will show that landmark genes exhibit both internal similarity within a landmark and external dissimilarity among landmarks.

#### Characteristics of landmarks hinting different cell types

It is evident that distinguishing different cell types requires a variety of distinct and informative features. To this end, we analyzed the information contained of different landmarks. Therefore, we randomly chose five landmarks constructed in LC_2_ (column vectors in *Q*_2_ matrix) each of them will capture a specific cell type, and show their non-relationship by calculating their Pearson correlation coefficients. Here, Camp2 and Romanov datasets are used in this analysis. As shown in Fig. 4a, the landmarks that capture different cell types are quite dissimilar and generally negatively correlated or irrelevant in the two datasets. This demonstrates that different landmarks can capture relatively typical information for different cell types, and the information associated to different cell types are mutually exclusive. In addition, the expression patterns of the five landmarks are visualized in Fig. 4b and 4c, which obviously show that each landmark is highly expressed in one specific cell type and hardly expressed in other cell types.

**Figure 4.**
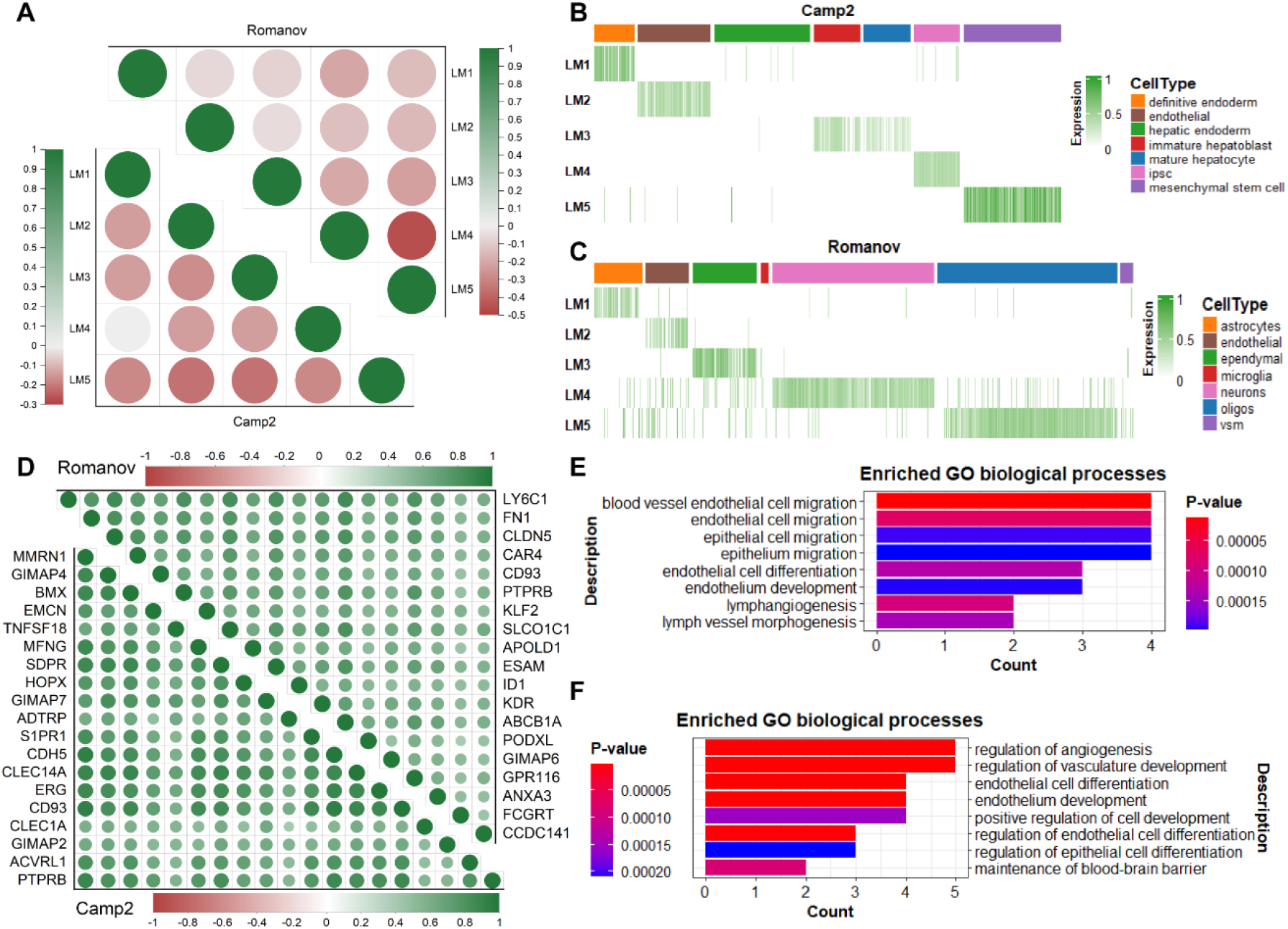
Charts of similarity of genes in the same landmark and dissimilarity between landmarks. (*A*) Dot plot of Pearson correlation coefficient of five landmarks identified by scQA on Camp2 and Romanov datasets. Similarities of landmarks in Camp2 dataset are showed in lower triangular matrix while upper triangular matrix shows similarities of landmarks in Romanov dataset, (*B*) Heatmaps of values of five landmarks in Camp2 datasets. Color bars on the top of heatmaps indicate the cell types. (*C*) Heatmaps of values of five landmarks in Romanov datasets. Color bars on the top of heatmaps indicate the cell types. (*D*) Dot plot of Pearson correlation coefficient of genes in one landmark identified by scQA on Camp2 and Romanov datasets. Landmark genes of Camp2 dataset are showed in lower triangular matrix while upper triangular matrix shows landmark genes of Romanov dataset. (*E*) GO biological processes (BP) enriched in genes in the same landmark of Camp2 datasets. Top eight terms are showed on the *y*-axis, and *x*-axis indicates the number of genes enriched. Color bar indicates the p-values. (*F*) GO biological processes (BP) enriched in genes in the same landmark of Romanov datasets. Top eight terms are showed on the *y*-axis, and *x*-axis indicates the number of genes enriched. Color bar indicates the p-values.

#### Characteristics of genes within same landmark

Since we have analyzed the significant differences in information captured by landmarks hinting different cell types, we next analyze whether genes of same landmark share similar expression patterns. To test whether scQA can combine genes with high consistency, we calculated the Pearson correlation coefficients of genes from same landmark (Fig. 4d, Supplemental Fig. S5, Fig. S6). To better display, here we used landmarks (LM2 in Fig. 4b and Fig. 4c) which correspond to the endothelia cell type that both contained in Camp2 and Romanov datasets for subsequent analysis, and other cell types can be found in Supplemental Fig. S5 and Fig. S6. As shown in Fig. 4d, upper triangular matrix showed the Pearson correlation coefficients between genes of endothelia cell type constructed in Romanov dataset, while lower triangular matrix was for Camp2 dataset. In addition, the number of genes in the two landmarks are both 19.

As we can see, all genes are positively correlated with each other and the correlation coefficients are above 0.5 for almost all similarities between genes located in LM2 of the two datasets. To further analyze the characteristic of genes in the same landmark, the expression landscape of all genes in LM2 for Camp2 and Romanov datasets were displayed in Supplemental Fig. S7. It can be observed that genes assigned to same landmark possess highly similar expression patterns. Accordingly, scQA can aggregate highly similar and informative genes in terms of expression values, and that these genes do facilitate cell clustering. Now we have seen that genes within one landmark have remarkably similar expression patterns, we proceeded to examine their biological similarity. To this end, we performed functional enrichment analysis using Gene Ontology (GO) biological processes on the gene sets of two landmarks analyzed above. We then selected top 8 terms from each gene set according to their p-value. As shown in Fig. 4e, for Camp2 dataset, the enriched GO biological processes of landmark genes in LM2 are “blood vessel endothelial cell migration”, “endothelial cell migration”, “endothelial cell differentiation”, “endothelium development” and other terms, which are directly related to the development of endothelial cells. Endothelial cells lie in the interior surface of blood vessels and lymphatic vessels, which means that genes associated with endothelium cells can indeed be relevant with the term “lymph vessel morphogenesis”. For Romanov dataset, the enriched biological processes are partially overlapping with Camp2 dataset, including “endothelial cell differentiation” and “endothelium development”. The endothelial cells are also involved in the formation of new blood vessels which referred to as angiogenesis. So genes in LM2 of Romanov dataset are also enriched in the functions related to endothelium, such as “regulation of angiogenesis” and “regulation of vasculature development”. Taken together, genes selected by scQA are of significant biological implications and interpretability. In terms of cell clustering, a landmark is formed by the combination of a set of similar genes, and scQA works by constructing multiple different landmarks to grab more valuable information to assist in clustering cells more precisely.

### External validation of landmarks

To further demonstrate the importance of the roles played by landmarks in cell clustering, we evaluate the reliability of landmark with external information. Our analyses covered the following three perspectives: association of landmark genes and differentially expressed genes (DEGs), hub genes and marker genes, and eight datasets are involved in this section, including Klein, Romanov, Segerstolpe, Zeisel, Camp1, Lawlor, Xin and Shekhar datasets.

#### Landmark genes reliably overlap with DEGs

To detect whether the selected landmark genes are informative genes, we conducted a series of comparative experiments by comparing the differentially expressed genes and landmark genes identified by scQA on Klein, Romanov, Segerstolpe and Zeisel datasets. To derive DEGs, we performed Wilcoxon Rank-Sum test implemented in the Seurat package along with Benjamini-Hochberg correction for adjusting p-value on four datasets. Firstly, for convenience, we use M1 to represent the set of DEGs where cells are assigned reference labels, M2 represents the set of DEGs where labels of cells are generated by scQA, and M3 is the set of landmark genes derived in LC_2_. Here, the genes with the absolute value of logFC above 2 and adjusted p-value less than 0.01 were selected as DEGs. As shown in Fig. 5, the three sets have a high degree of overlap in the Venn diagram. For Klein dataset (Fig. 5a), the overlap odd (Supplemental Method 3) of M1 and M2 of DEGs achieved 99.2% (34.3% + 64.9%), and M3 of landmark genes was wholly contained in the intersection of M1 and M2. For Romanov dataset (Fig. 5b), M3 was wholly encompassed in the union of M1 and M2 and the overlap odd of M1 and M3 achieved 93.9%. For Segerstolpe and Zeisel datasets (Fig. 5c and Fig. 5d), we can notice that extremely few points from set M3 of landmark genes were absent in the intersection of sets M1and M2 of DEGs, with 3 genes being in M3 but not in the intersection of M1 and M2 (one gene was not in M1, one gene was not in M2 and the other was not in the union of M1 and M2) for Segerstolpe dataset and 29 genes being in M3 but not in the intersection of M1 and M2 for Zeisel dataset. The two sets, M1 and M2 nearly coincide exactly indicating that our clustering results are fairly reliable compared with the reference labels. On the other hand, the virtually complete inclusion of M3 in M1 reflects the fact that our feature extraction strategy is quite reliable, but may drop partial information by performing the low-dimensional representation. Since M1, M2 and M3 are highly identical, we conducted Wilcoxon Rank-Sum test on set M3 of landmark genes identified by scQA to show the ability of scQA for selecting informative genes on Klein and Zeisel datasets (Fig. 5e, Supplemental Fig. S8), four genes displayed in Fig. 5e and nine genes plotted in Supplemental Fig. S8 are the top ranked DEGs in accordance with adjusted p-value in each cell type on Klein and Zeisel datasets respectively. It is obvious in Fig. 5e, *FAM25C* and *KRT18* are up-regulated genes in the cell development of day 0 and day 7 respectively while other two genes, *AHNAK* and *RN28S1*, are down-regulated in the cell development of day 2 and day 4 in Klein dataset. In addition to this, Supplemental Fig. S8 showed that some landmark genes are label-specific, like *DLX6OS1, GM11549, ELTD1, ACSBG1, MSX1, MYL9*, which are only expressed in some specific cell type. And other genes, like *HPCA*, exhibiting different expression patterns in multiple cell types which is highly expressed in CA1 pyramidal cells, lower expressed in interneurons, S1 pyramidal cells and barely expressed in other types of cells. This phenomenon implies that genes which are characterized with binary expression patterns indeed recognize some clusters, but quantitative expression patterns of genes are of importance for further distinction. One cannot be without the other for deriving precise clusters.

**Figure 5.**
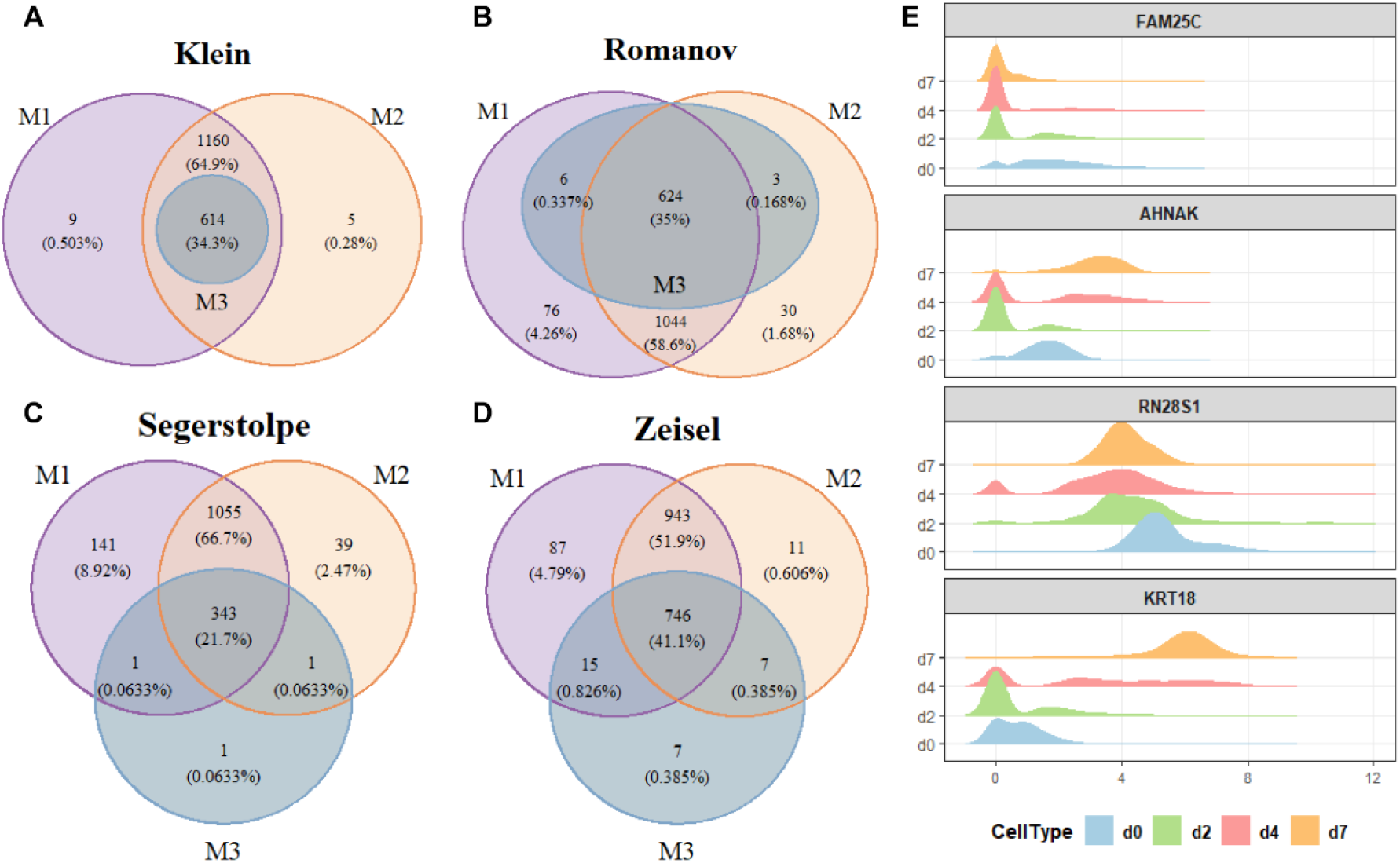
Comparison of DEGs and landmark genes identified by scQA. (*A-D*) Venn diagrams of M1, M2 and M3 on Klein, Romanov, Segerstolep and Zeisel datasets, respectively. M1 is the set of DEGs which Wilcoxon Rank-Sum test is conducted in terms of the reference labels, M2 is the set of DEGs with labels of cells from cluster results of scQA, and M3 is the set of landmark genes identified in scQA. (*E*) Joy plot of feature genes extracted by scQA on Klein dataset with landmark genes marked on the title of each subgraph, expression values represented on the *x*-axis and color representing the cell types.

#### Landmark genes are generally hub gene

As weighted gene co-expression network analysis (WGCNA) can be used to detect highly correlated genes and genes with strong connections in expression network, which are regarded as hub genes as well as considered functionally significant. We identified hub genes from different datasets which significantly overlapped with the landmark genes generated by scQA. As shown in Table 1, the number of hub genes were set to 100 and the number of landmark genes respectively in Camp1, Lawlor, Segerstolpe and Xin datasets and intersection of hub genes and landmark genes were listed. In Camp1 dataset, 100 out of 2,000 genes are regarded as hub genes, and 99 out of 100 hub genes can be found in the 464 landmark genes identified by scQA. Then, we enlarged the number of hub genes to the number of landmark genes, and 75% (348/464) hub genes are identified by scQA. Similar results can be observed in the other three datasets. To measure such significance, we conducted a hypergeometric test (Supplemental Method 4) to estimate the p-value with correction based on the results of randomized trials. The results showed that our method is extremely significant for selecting landmark genes being hub genes, as the adjusted p-value is much less than 0.01 which is considered to be quite significant on both cases of selecting hub genes.

**Table 1.**
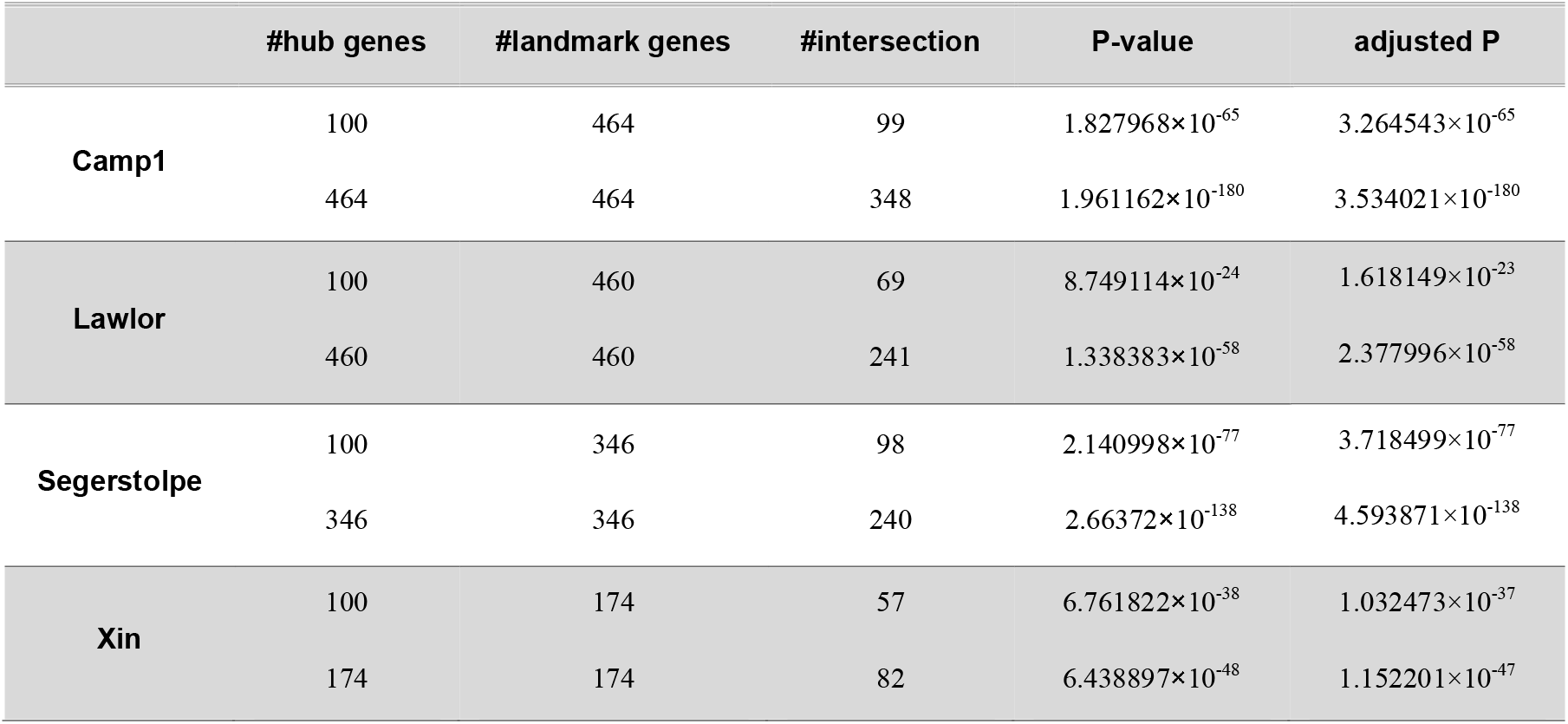
Comparison of hub genes and feature genes identified by scQA in Camp1, Lawlor, Segerstolpe and Xin datasets.

#### Landmark genes comprise marker genes

In the following, we analyze the reliability of landmark genes by comparing them with marker genes. The marker genes were downloaded from Panglao DB (https://panglaodb.se/). Since we pre-processed the expression matrix by selecting 2,000 genes for clustering analysis, the subsequent analyses were performed on these 2,000 genes on Xin and Romanov dataset. We use HVGs to denote the set of 2,000 highly variable genes. LMGs represents the landmark genes, while the marker genes are split into two sets: MKG1, which represents marker genes in the set of landmark genes, and MKG2, which represents marker genes not in the set of landmark genes. As shown in Fig. 6a, the sum of MKG1 and MKG2 represents the number of marker genes contained in 2,000 genes, which accounts for 2.1% of 2,000 genes and specifically 42 genes for Xin dataset. In addition, 174 genes are selected as landmark genes using scQA, which cover 22 marker genes. We then investigated if scQA is effective in enrolling marker genes of all cell types. The analysis revealed that over 60% of marker genes associated with alpha cells and beta cells were found in landmark genes, along with 40% of marker genes related to gamma cells, and 30.8% marker genes of delta cell type, which had the lowest percentage of marker genes found in scQA (Fig. 6a, right). We further conducted a hypergeometric test to evaluate the significance of this result, and found it to be extremely significant in three of four cell types (alpha, beta and gamma) with p-values less than 0.01. In details, the p-values are presented in Supplemental Table S7. We also analyzed the Romanov dataset that include 7 cell types, as shown in Fig. 6b. From the chart we can see a remarkable difference number of marker genes of different cell types contained in landmark genes, varying from 74.4% to 6.7%. One possible factor causing this phenomenon is the great variation in the number of cells of different cell types in the dataset. In particular, the cell type with the lowest percentage of marker genes in landmark genes tends to be the cell type with the lowest number of cells in the dataset. For instance, in the Romanov dataset, only 48 cells of microglia are included, which is 1.6% of all 2,881 cells. Consequently, the expression values of genes expressed in only 1.6 percent of cells are plausibly dampened as random expression values or noises of no significance.

**Figure 6.**
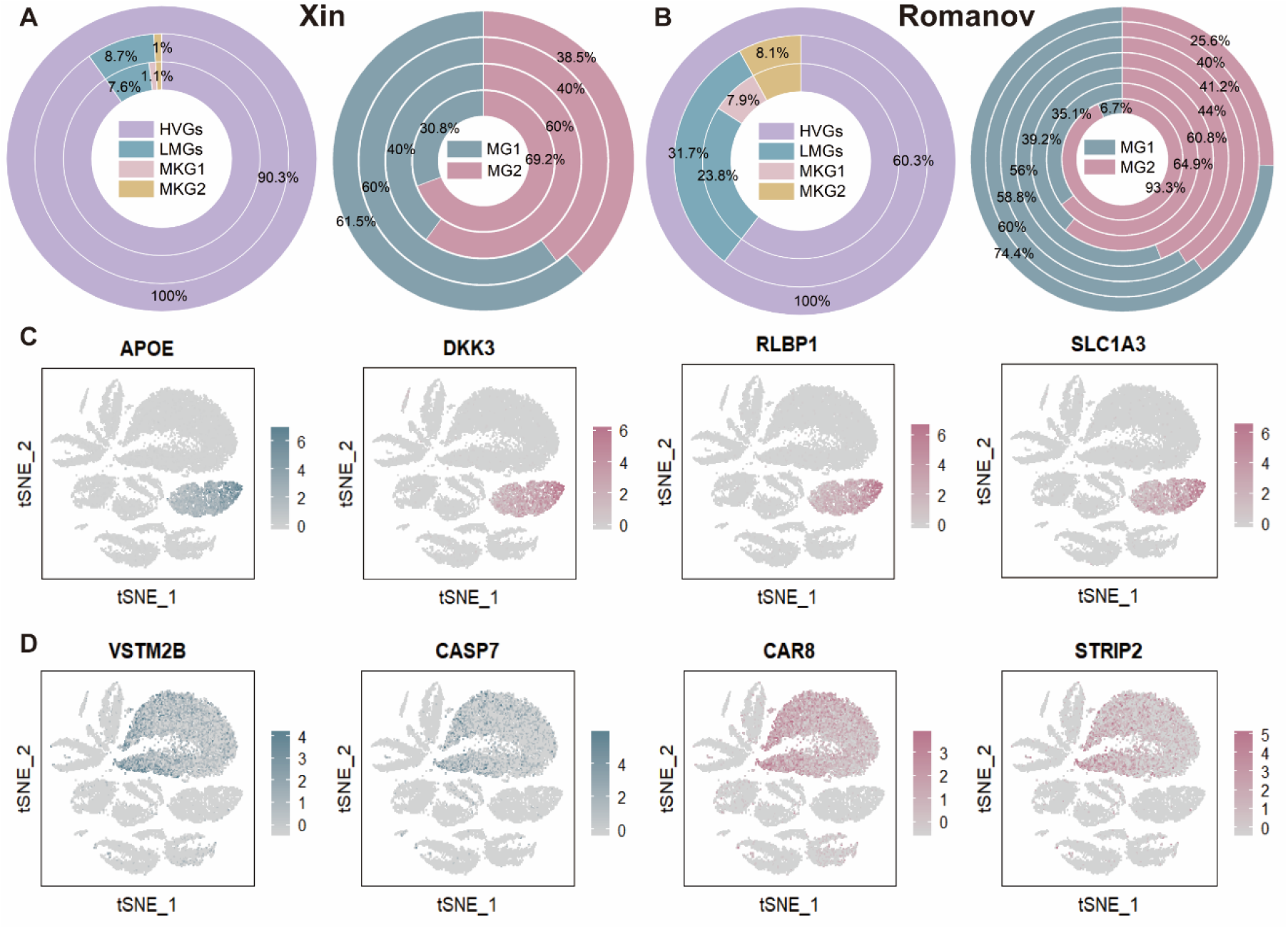
Comparison of marker genes and landmark genes identified by scQA. (*A*, left) Doughnut chart of percentage of landmark genes and marker genes in Xin dataset. (*A*, right) Doughnut chart of percentage of marker genes in and not in LMGs denoted by MG1 and MG2 respectively of four cell types in Xin dataset. Rings refer to alpha, beta, gamma and delta cell types from the outer to the inner, (*B*, left) Doughnut chart of percentage of landmark genes and marker genes in Romanov dataset, (*B*, right) Doughnut chart of percentage of marker genes in and not in LMGs denoted by MG1 and MG2 respectively of seven cell types in Romanov dataset. Rings refer to ependymal cells, vascular smooth muscle cells, oligodendrocytes, neurons, endothelial cells, astrocytes and microglia from the outer to the inner. (*C*) t-SNE visualizations of genes related to müller glia cell type in Shekhar dataset. (*D*) t-SNE visualizations of genes related to rod bipolar cell type in Shekhar dataset.

Additionally, due to the incomplete record of marker genes in the database, we examined genes that belong to the same landmark group as the marker genes recorded in the original literature of the Shekhar dataset. The Shekhar dataset contains five major cell types: müller glia (MG), rod photoreceptors, cone photoreceptors, amacrine and bipolar cells (Supplemental Fig. S9), with the latter further subdivided into 14 subtypes (Supplemental Fig. S10). We focused our analysis on one major cell type (MG) and one subtype of BC. *APOE* is a major markers of MG cells in the retinal class and was used in the original literature (Shekhar et al. 2016). Therefore, we extracted some genes from the landmark group that contains *APOE*. As illustrated in Fig. 6c, all of these genes, including *APOE, DKK3, RLBP1* and *SLC1A3*, exhibit similar expression patterns that are expressed almost exclusively in MG cells. The gene *APOE* exhibits a striking ability to express in MG cells with non-zero expression values observed in all MG cells, but also observed in other cell types with other cell types expressed accounting for 51.8 percent of all expressed cells (Supplemental Table S8). The other three genes appear to exhibit more specificity with 74.8%, 74.2% and 83.9% of expressed MG cells in all expressed cells respectively. Unfortunately, they are not expressed in all MG cells with an expression percentage of more than 96. Subsequently, we found that the three genes were also used as marker genes of MG cells in several literatures (Macosko et al. 2015; Liang et al. 2019; Gautam et al. 2021). This indicates that scQA can excavate similar marker genes form the landmark, which contributes a lot for cell type identification. Next, we considered another cell type, rod bipolar cells (RBCs), which is a subtype of bipolar cells with relatively large number of cells (10,888 cells of 26,830 cells in Shekhar dataset). In the original study, two genes, *VSTM2B* and *CASP7*, were identified as related to RBCs. We investigated the landmark group encompassing these two genes and identified multiple genes that showed similar expression patterns (Fig. 6d, Supplemental Fig. S11). These genes are expressed in over 80% of cells being RBCs. However, since the number of RBCs is quite large, not all RBCs expressed in these genes. Our analysis showed that *CAR8* manifests a higher percentage of being expressed comparing with *VSTM2B* and *CASP7*, with a percentage of 85.1% (Supplemental Table S8), which has been confirmed as a marker gene in previous studies (Kim et al. 2008; Puthussery et al. 2011). Another gene, *STRIP2*, also exhibits strong specificity in expression with a percentage of 90.7% of expressed cells being RBCs which we have not found in any reference recorded as a marker gene. *STRIP2* may be relevant to retinal development since a recent study highlighted the importance of *STRIP1* in inner retina development (Ahmed et al. 2022) and both *STRIP1* and *STRIP2* play a role in regulating cell morphology and cytoskeletal organization and are paralogs to each other. Thus, scQA has the potential to detect new marker genes.

## Discussion

For the past decade, researchers have been working on analyzing single-cell RNA-seq datasets. One of the most challenges is to tackle dropouts appropriately. Increasing evidences have shown that dropouts should not be discarded at the beginning of the downstream analysis. It has been observed that the dropout events are mainly inherited from biological side instead of technical side for UMI-based data. Once cells are correctly grouped according to their heterogeneity, most dropouts will disappear. However, imputation or other preprocessing methods, which attempt to remove dropouts, may inevitably introduce biases. In this study, we introduce a new algorithm to fully integrate information from both dropouts and quantitative gene expressions of different levels. Specifically, we first identify informative genes as chunks of genes (landmarks) exhibiting similar expression trend. To reduce the impacts of orders of genes on clustering results and save running time, we offer some predefined clusters consisting of genes with high consistency as prior information. LC is designed for effectively aggregating genes with similar qualitative expression patterns or quantitative expression patterns to ensure that landmark genes exhibit internal similarity and external dissimilarity, and selected genes turn out to be informative since we demonstrated that there is a high degree of overlap with landmark genes and differential expression genes associated with true labels. In addition to this, we illustrated landmark genes were generally hub genes derived by constructing weighted gene co-expression network which have been considered functionally significant. We also demonstrated that landmark genes comprise marker genes and scQA has the potential to detect new marker genes to some extent.

Once a landmark being obtained, all genes from the landmark are corrected to each other to refine themselves. Finally, qualitative and quantitative landmarks are regarded as features to be fed into CC module, where cell clusters will be formed. A novel label propagation strategy is adopted in CC by performing seed generation, label propagation, pruning and merging and cluster expansion. Instead of assigning a unique label to each cell, we only grant seeds labels, in contrast to traditional label propagation algorithm (Raghavan et al. 2007). And the threshold of seeds is determined by the pre-grouping of cells in accordance with qualitative landmarks. After defining the labels of seeds, we require to propagate the labels by considering both neighbors of seeds and with seeds as neighbors. We also incorporate an alert mechanism to prevent cells from being wrongly labeled in the absence of enough information. After pruning and merging clusters, all retained clusters are expanded based on a newly defined indicator, and consistency clustering results are obtained through iterative label propagate process. The results show that scQA substantially outperforms other popular methods.

scQA is a reliable automated clustering tool for single-cell RNA-seq datasets with various number of cells, but limitations still stand in scQA. In the pre-processing step, only 2,000 HVGs are retained to perform subsequent clustering analysis resulted in some informative genes being discarded (Supplemental Table S9). Therefore, the inclusion of priori information to retain more informative genes can be an option. Since clusters are subject to pruning and merging, rare cell types may be left out and two sub-types may be recategorized into one cluster. Similar to other clustering methods applying similarities or distance metrics, genes are vulnerable to misclassified as the impact of the high dimensionality and similarities are required for constructing initial clusters in LC. In the future, we will optimize the process of selecting HVGs, and adapt scQA to multi-omics data since it has a natural structure that can integrate two types of information.

## Methods

### Data pre-processing

To reduce the sparsity of original scRNA-seq data, we first filter genes expressed in less than *x*% cells and those expressed in more than (100 – *x*)% cells. The filtered genes are considered to be less informative. The parameter *x* is set to 2.5 in our experiments. For further picking out information-rich genes, we retain 2,000 highly variable genes according to the method in (Brennecke et al. 2013; Satija et al. 2015). All the data are log-transformed. The resulting matrix is then normalized such that every entry in the matrix falls in [0, 1]. We by *D_m×n_* denote the preprocessed gene expression matrix, where *m*=2,000 corresponds to the informative genes selected in the data pre-processing step, and *n* the number of cells. The entry *d_ij_* in *D* represents the expression value of gene *i* in cell *j*.

### Qualitative Landmark constructor

LC_1_ module is in charge of generating landmarks to form low-dimensional representation in binary form. The pseudocode for this process is shown in Agorithm1.

For each of these genes in *D*, we define a binary vector *G_i_* = (*g_i1_*, *g_i2_*, …, *g_in_*). If *d_ij_* > 0, *g_ij_* = 1; otherwise, *g_ij_* = 0. In order to reduce the complexity of computation and to increase the stability of clustering, we first sort the genes in order of decreasing number of zeros in each gene vector. For each gene *G_i_*, we take 0.05*m*. (=100) genes respectively in front of and behind *G_i_* in the sorted sequence as the neighborhood of *G_i_*, and calculate the similarities between the gene *G_i_* and all genes in the neighborhood of *G_i_* as(*n* —(*G_i_, G_j_*))/*n*, where *h*(*G_i_, G_j_*) is Hamming distance between *G_i_* and *G_i_*. We then construct gene similarity graph with vertices representing genes and edges the top 1,999 gene pairs (one thousandth of all gene pairs) in similarity scores. Then connected components with more than two vertices in the graph form the initial gene clusters *C_r_* (*r* = 1,2, … ). To assign genes outside the current gene clusters, we, for each cluster *C_r_*, define a template vector *T_r_* = (*t_r1_*, *t_r2_*, … *t_rn_*), which is obtained by rounding the mean of vectors of the genes in the cluster, i.e., *t_rk_* = 1 if ∑_*G_j_*∈*C_r_*_ *g_jk_*/|*C_r_*|≥0.5, and 0 otherwise. Obviously, if a gene outside union of current clusters is of its binary expression vector exactly same with one of the current template vectors, then the gene should be assigned to the cluster of the template vector. To assign the genes whose binary expression vector differing from any template vectors, we found a way to accurately and reliably define the similarity score between a gene *G_i_* that has not been clustered yet and a current cluster *C_j_*. By *μ_ijk_* we denote the similarity increment contributed by the gene *G_i_* and the cluster *C_j_* at the *k*th component. We may specify *μ_ijk_* = 1 if *g_ik_* = *t_Jk_*, 0 if *g_ik_* = 1 Λ *t_jk_* = 0, and *s_jk_* if *g_ik_* = 0 Λ *t_jk_* = 1, where *s_jk_* is *k*th component of a subsidiary vector *S_j_* defined for each current cluster *C_j_* as follows:

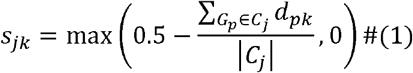

The similarity *s*(*G_i_*, *C_j_*) between the gene *G_i_* and the cluster *C_j_* is defined as average similarity increments contributed by *G_i_* and *C_j_* at all components, i.e., 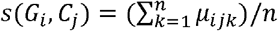. The rationality can be strongly convinced from the definition of similarity alone: the *k*th component contributes to the entire similarity a ceiling score 1 when *g_ik_* = *t_jk_* because *g_ik_* = *t_jk_* if and only if either *g_ik_* = *t_jk_* = 0 or *g_ik_* = *t_jk_* = 1. The *k*th component contributes to the entire similarity a null score 0 when *g_ik_* = 1 and *t_jk_* = 0 because this situation means that genes in current cluster *C_j_* do not express at the *k*th component while *G_i_* does. The *k*th component contributes to the entire similarity a tricky score *S_jk_* when *g_ik_* = 0 and *t_jk_* = 1 since this situation makes it a little tricky to reasonably assess the score that the *k*th component contributes. Notice that *g_ik_* = 0 may be from the fact that the expression of *G_i_* under *k*th cell was not observed due to its weakness, that is to say that *G_i_* with some chance may have real positive expression value but not prominent even in the case *g_ik_* = 0, and, the more prominent the expression the smaller the chance. We may assume that with no chance we have *d_ik_* > 0.5, and that with chance 0.5 – *x* we have *d_ik_* = *x* for *x* ∈ (0, 0.5]. This is the reason why we defined the score *S_jk_* contributed by *G_i_* and *C_j_* at *k*th component as that in (1).

For genes that have not been clustered yet, we iteratively calculate the similarity *s*(*G_i_*, *C_j_*) between gene *G_i_* and cluster *C_j_* (Algorithm 1, lines:8-10), and add *G_i_* to *C_j_* if *s*(*G_i_*, *C_j_*) exceeds a prespecified parameter *ρ*, and update the template vector *T_j_* and subsidiary vector *S_j_* accordingly (Algorithm 1, lines:12-14), or form a new singleton cluster (Algorithm 1, lines:14-16). After all genes clustered, we remove those clusters of less than 3 genes (Algorithm 1, lines: 18-20), and reduce the parameter *ρ* for another round iteration if necessary. By default, we iterate once with the fixed parameter *ρ* set to 0.7, and use all retained clusters to create a qualitative cell-landmark matrix *Q*_1_ with cells corresponding to rows and landmarks to columns which are consensus pattern of their corresponding clusters (Algorithm 1, line:23).

#### Algorithm 1: Construct Qualitative Landmarks

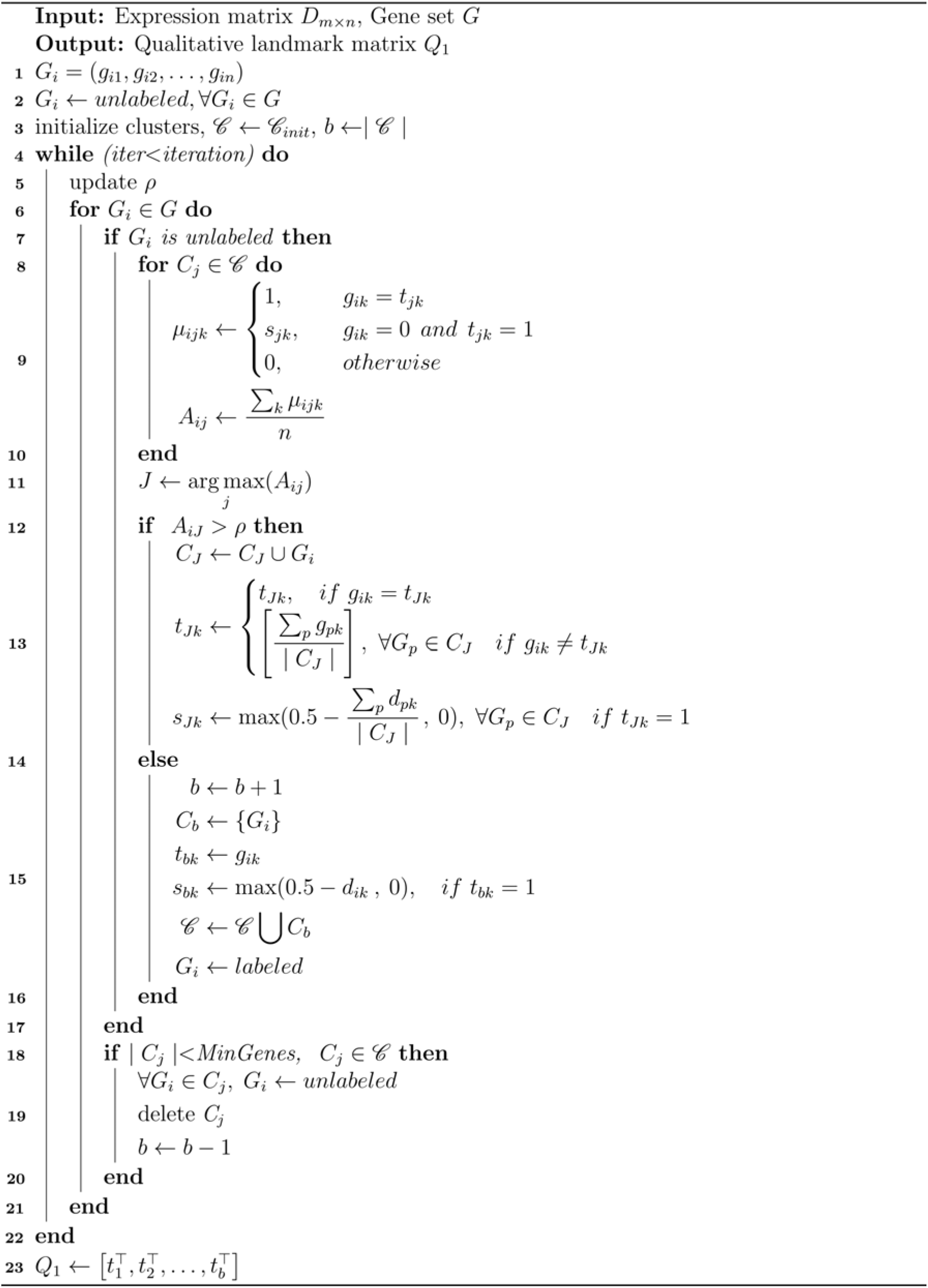

### Quantitative Landmark constructor

LC_2_ module is adopted to form low-dimensional representation of cells by generating a quantitative cell-landmark matrix with the pseudocode being shown in Algorithm 2 below. We first discretize the continuous expression values by binning. To do so, we divide the interval [0, 1] into *k* bins, [0, 0], (0, 1/(*k* – 1)], (1/(*k* – 1), 2/(*k* – 1)], …, ((*k* – 2)/ (*k* – 1), (*k* – 1)/(*k* – 1)], which are indexed by 1, 2, …, *k*, so that every expression value must falls in/on one of them since they have been normalized. The default value of *k* is set to 6 in our experiments. Then the gene expression matrix *D* can be qualitatively represented, denoted by *D’*, by replacing *d_ij_* with bin index *s* if and only if *d_ij_* falls in sth bin. After qualitative representation, we group genes in the number of bins they used, i.e., two qualitative represented genes belong to a single group if and only if they are binned using same number of bins. Similar to LC_1_, we create a quantitative cell-landmark matrix for each group and then merge them into a single one cell-landmark matrix. Doing so will greatly save in running time and get improved in accuracy.

#### Algorithm 2: Construct Quantitative Landmarks

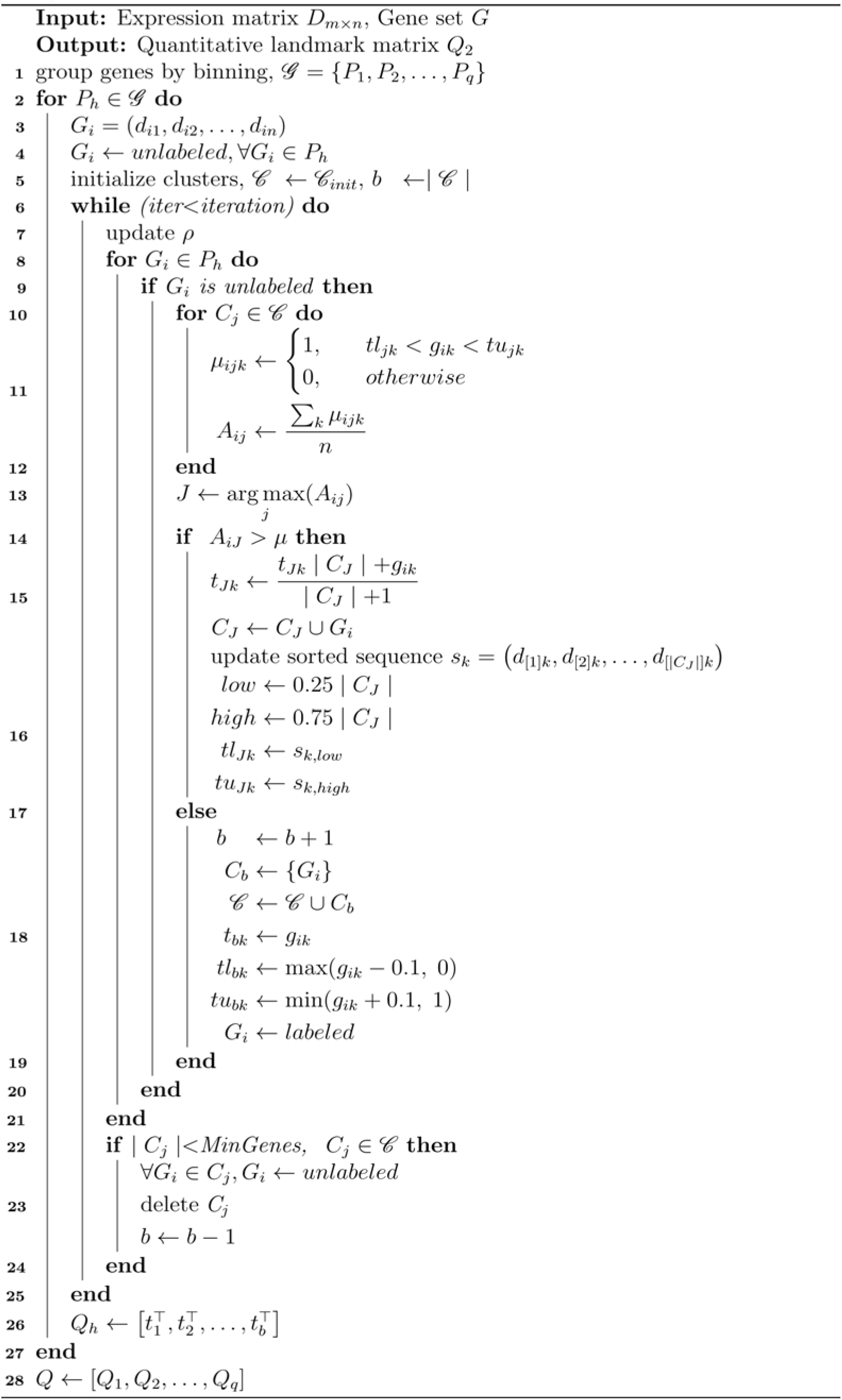

What differs to LC_1_ is that similarity scores are calculated based on Spearman’s rank correlation coefficient. After deriving initial clusters as did in LC?, we have also template vectors which are simply the mean vectors of the gene expression vectors in their respective clusters. For clustering genes of quasi-trend-preserved, we define upper and lower bound vectors *TU_j_* and *TL_j_* for each template vector *C_j_*. The lower (resp. upper) bound vector *TL_j_* (resp. *TU_j_*) with its *k*th component being the element at the lower (resp. upper) quartile *x*%|*C_j_*| ((1 – *x*)%|*C_j_*|) in an ordered sequence of |*C_j_*| elements in the *k*th component of cluster *C_j_*. For a gene *G_i_* outside clusters *C_j_*, we define their similarity *s*(*G_i_*, *C_j_*) as the ratio between the number of components where the expression values of *G_i_* fall between the lower and upper vectors and *n* (Algorithm 2, lines:10-12). We then add *G_i_* to *C_j_* if *s*(*G_i_*, *C_j_*) > *ρ* and update the template vector *T_j_*, the upper bound vector *TU_j_* and the lower bound vector *TL_j_* (Algorithm 2, lines:14-17), and create a new singleton cluster otherwise (Algorithm 2, lines:17-19). After all genes clustered, we remove those of less than 3 genes (Algorithm 2, lines:22-24), and reduce the parameter *ρ* for another round iteration if necessary. By default, we iterate once with a fixed threshold 0.7, and all retained clusters of all groups are jointed to create a quantitative cell-landmark matrix *Q*_2_ with cells corresponding to rows and landmarks to columns which are consensus of the clusters (Algorithm 2, line:26).

### Cluster constructor

CC module clusters cells by combination of the two cell-landmark matrices respectively obtained in LC_1_ and LC_2_ modules with the pseudocodes shown in Algorithm 3. We construct a directed nearest neighbor graph *G* with nodes representing cells, and edges representing cell pairs of higher similarity obtained using Pearson correlation coefficient based on quantitative cell-landmark matrix to avoid bias from use of the same similarity metric as in landmark constructor. For each node in *G*, it has 0.1*n* (resp. 0.05*n*) out neighbors if *n* < 10,000 (resp. *n* ≥ 10,000). The CC module is implemented in four steps:

#### Algorithm 3: Cluster Cells

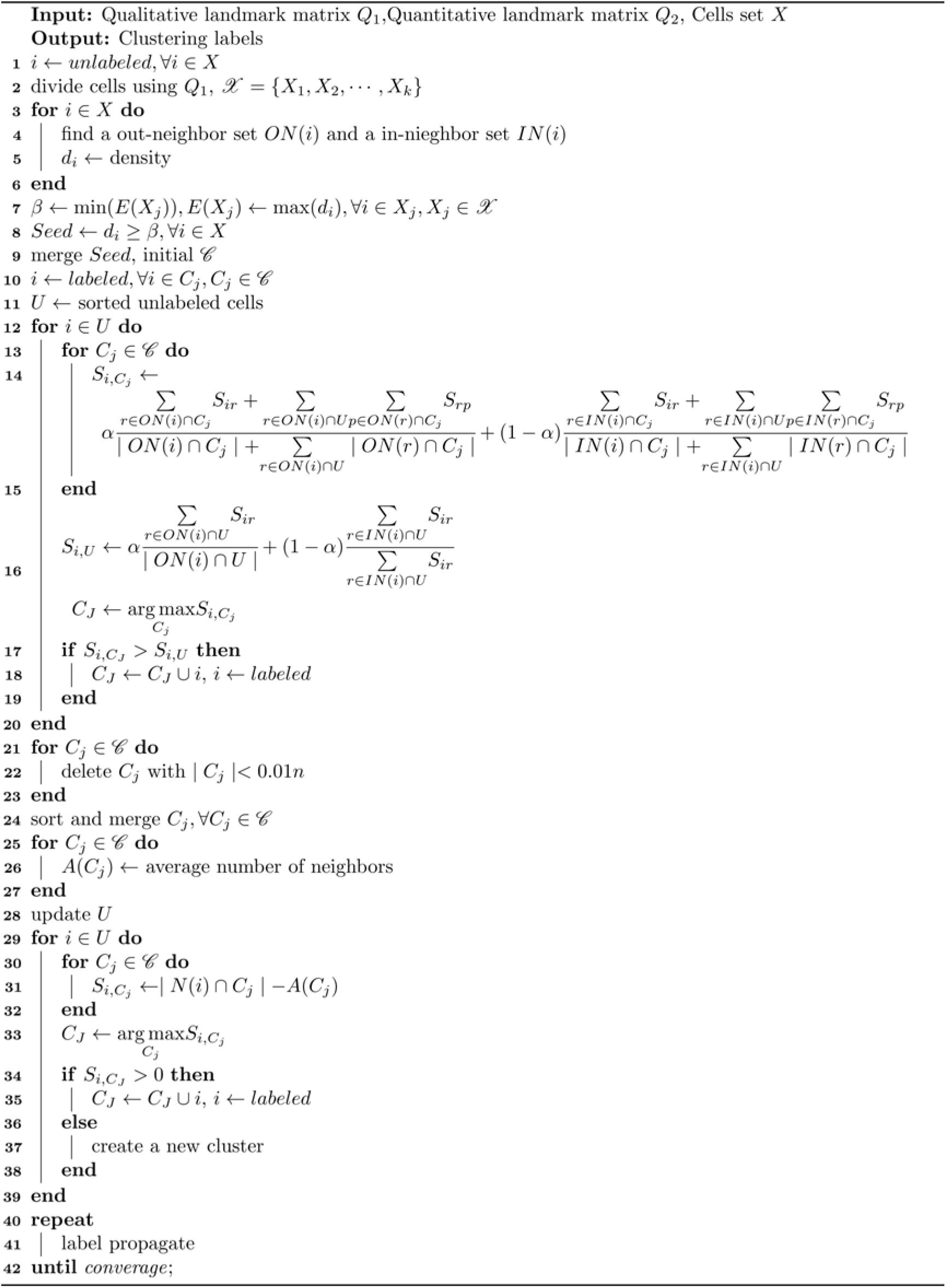

#### Seed generation

We first group cells based on the qualitative cell-landmark matrix *Q*_1_ such that cells in same group exhibit exact the same row vector in *Q*_1_. One of ideas behind scQA is that cells within a same group should form a core of a final cluster to be tested. That being the case, we can generate all the potential seeds following the guidance of the core groups. Doing so, we assign to each node a density *d* which defined as the average of the similarities between the node and all its neighbors. Intuitively, the higher the density of a node, the higher the possibility of the node being a real seed. We now determine a threshold *β* that a node of its density beyond should be qualified to be a reliable seed. To do so, we calculate the density for each node and record the maximum one for each group. Then the minimum one among the maximum densities each is from a group is used as the threshold *β*, i.e., nodes of their densities larger than this threshold are used as seed candidates (Algorithm 3, lines:7-8). If two seed candidates that are mutually neighbors in *G*, we merge them as single one, and assign to each seed a unique label. We then treat every seed as an initial cluster. All other nodes are sorted in a decreasing order of their densities.

#### Label Propagation

For each node *n_i_* outside the current clusters *C_j_*, we calculate the similarity *S_iC_j__* between the node *n_i_* and each current cluster *C_j_*, i.e.,

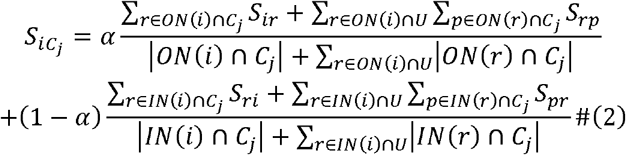

Where *ON*(*i*) represents the set of out-neighbors of the node *n_i_*, *IN*(*i*) the set of in-neighbors of the node *n_i_*, *U* the set of unlabeled nodes, and *S_ij_* the similarity between nodes *n_i_* and *n_j_*. The first term on the right side of the equation can be viewed as the similarity from node *n_i_* to cluster *C_j_* obtained via dividing the sum of similarities on the paths from *n_j_* to each node in *C_j_* at distance at most two by the number of these paths. The second term on the right side of the equation can be viewed as the similarity from cluster *C_j_* to node *n_i_* obtained via dividing the sum of similarities on the paths from each node in *C_j_* to the node *n_i_* at distance at most two by the number of these paths. The *α* is a combinatorial coefficient with default set to 0.5 (Algorithm 3, lines:13-15). For a set *U* of unlabeled nodes, we by *S_iU_* denote the average similarity between *n_i_* and *U* obtained via dividing sum of similarities on all edges from *n_i_* to *U* or from *U* to *n_i_* by the number of these edges (Algorithm 3, line: 16). We now let *U* be the whole unlabeled nodes without *n_i_*, and 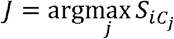. If *S_iC_j__* > *S_iU_*, we add the node *n_i_* to the cluster *C_j_*, and label it; otherwise, we do not believe there is enough information to label this node and the next node in *U* will be considered.

Repeat the procedure until all nodes are processed in *U*.

#### Pruning and Merging

Pruning is to remove those too small clusters. After removing, we prioritize merging those retained smaller ones such that similar ones are merged into a single one (Algorithm 3, lines:21-23). By *S_ij_*, we define the similarity between cluster *C_i_* and cluster *C_j_*, which is equal to the number of edges from cluster *C_i_* to cluster *C_j_*. Clusters are sorted by size, and if the maximum similarity between the current cluster and larger clusters, denoted by 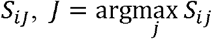, exceeds number of edges within the cluster *C_i_*, cluster *C_i_* will be merged into cluster *C_j_*. Clusters who have absorbed other clusters cannot be merged into a larger cluster.

#### Cluster Expansion

For each retained cluster, we by *A* denote the average number of neighbors in the cluster of nodes of the cluster. For each unlabeled node, we redefine the similarity between the node and a cluster as the difference between the number of neighbors in the cluster of the node and the *A* of the cluster (Algorithm 3, lines:30-32). The larger the difference is, the greater the possibility of the cell belonging to the cluster. If all the differences are less than 0, we may assume that this node should not be added to any current clusters, and instead, we create a new cluster in which this node resides. So far, all cells are labeled, we iterate the label propagation strategy until converge.

### Comparison with other tools

As mentioned before, SC3 is a consensus method developed by performing *k*-means on multiple matrices. Seurat is developed by performing PCA followed by recalculating the similarities based on shared nearest neighbors. CIDR is an ultrafast method developed by performing PCA based on zero-imputed similarities. Like SC3, pcaReduce is an agglomerative clustering approach developed by combining PCA and *k*-means. SIMLR is developed by learning a similarity matrix via combining multiple kernels. TSCAN is designed by combining PCA with model-based clustering method. RaceID2 is designed based on Pearson correlation coefficient to identify rare and abundant cell types using *k*-medoids. scGNN is a deep learning-based method and integrates graph convolutional network (GCN) into multi-autoencoder to improve clustering results. Based on Fuzzy C Mean (FCM) and Gath-Geva algorithms, scHFC performs a hybrid fuzzy clustering. For SC3 method, a hybrid approach combining SVM was utilized when cells in the dataset are more than 5,000. When cells exceed 10,000 in the dataset, large scale SIMLR was used as a substitution. scGNN was performed using LTMG. The range of estimation of *k* was set to 2 to 50 if it was needed to be specified. For the other parameters used in these methods, their default values were used in our comparisons in this paper.

### Performance evaluation

To measure the accuracy of the algorithm, we adopted four commonly used metrices: Adjusted Rand Index (ARI), Normalized Mutual Information (NMI), Fowlkes-Mallows Index (FMI) and Jaccard Index (JI). Certainly, the arrangement of cluster labels values will not change the score in any metric. The four indexes are calculated between the set *I* of identified cell clusters and the set *A* of annotated cell groups. Let *a* be the number of pairs of cells that are both in the same cluster in *I* and in the same group in *A; b* be the number of pairs of cells that are in the same group in *A* but in different clusters in *I*, *c* be the number of pairs of cells that are in different groups in *A* but in the same cluster in *I*, and *d* be the number of pairs of cells that are in different groups in both *A* and *I*. The four indexes are respectively defined as:

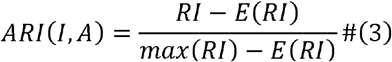

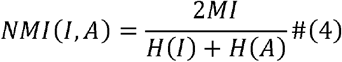

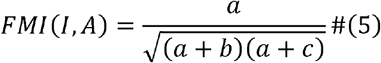

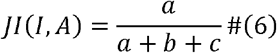

where *RI* is Rand Index of *I* and *A* defined by 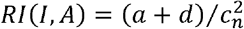. Entropy *H* is the amount of uncertainty defined by 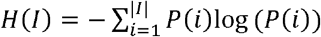, where 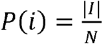. The mutual information (MI) between *I* and *A* is calculated by 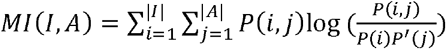.

## Data availability

Twenty publicly available scRNA-seq datasets were collected from https://hemberg-lab.github.io/scRNA.seq.datasets/, https://www.ncbi.nlm.nih.gov/geo/, https://www.ebi.ac.uk/ and R package “scRNAseq” for performance evaluation. The accession numbers of all datasets are: Biase dataset (Biase et al. 2014)(GSE57249), Yan dataset (Yan et al. 2013)(GSE36552), Deng dataset (Deng et al. 2014)(GSE45719), Camp1 dataset (Camp et al. 2015) (GSE75140), Lawlor dataset (Lawlor et al. 2017)(GSE86469), Camp2 dataset (Camp et al. 2017)(GSE81252), Xin dataset (Xin et al. 2016)(GSE81608), Baron1 dataset (Baron et al. 2016)(GSE84133), Muraro dataset (Muraro et al. 2016) (GSE85241), Segerstolpe dataset (Segerstolpe et al. 2016)(E-MTAB-5061), Hermann dataset (Hermann et al. 2018)(GSE109033), Klein dataset (Klein et al. 2015)(GSE65525), Romanov dataset (Romanov et al. 2017)(GSE74672), Zeisel dataset (Zeisel et al. 2015)(GSE60561), Baron2 dataset(GSE84133), Aztekin dataset (Aztekin et al. 2019)(E-MTAB-7716), Chen dataset (Chen et al. 2017)(GSE87544), Zilionis dataset (Zilionis et al. 2019)(GSE127465), Campbell dataset (Campbell et al. 2017)(GSE93374) and Shekhar dataset (Shekhar et al. 2016)(GSE81904). Details about the datasets are listed in Supplemental Table S2.

## Software availability

All source codes are publicly available at GitHub (https://github.com/LD-Lyndee/scQA).

## Acknowledgements

This work was supported by National Science Foundation of China with codes 11931008 and by Ministry of Science and Technology of China with code 2020YFA0712400.

